# RNF25 restrains GCN2 hyperactivation to sustain protein synthesis and cell proliferation in response to RNA damage

**DOI:** 10.64898/2026.03.21.713335

**Authors:** Antonia S. Seidel, Lucia Nemcekova, Martin Grønbæk-Thygesen, Xinyao Shi, Sofia Ramalho, Kelly C. Mordente, Simon Bekker-Jensen, Peter Haahr

**Affiliations:** Center for Gene Expression, Department of Cellular and Molecular Medicine, Faculty of Health, University of Copenhagen, Blegdamsvej 3B, 2200 Copenhagen N, Denmark

## Abstract

Regulation of protein synthesis is essential for maintaining cellular homeostasis during stress. The integrated stress response (ISR) is a conserved signaling pathway that modulates global mRNA translation through four eIF2α kinases—GCN2, PKR, PERK, and HRI. However, how these kinases are selectively activated and tuned to distinct stress signals to direct appropriate cell fate decisions remains poorly understood. Here, we employ ultra-deep mutagenesis screens to systematically map regulators of protein synthesis across diverse stress perturbations in human cells. This comparative approach identifies stress-specific translational control factors, including a previously unrecognized role for the E3 ubiquitin ligase RNF25 in selectively sustaining translation following UV irradiation and other RNA-damaging treatments. In this context, we demonstrate that RNF25 operates independently of its partner RNF14, and that its ubiquitin ligase activity, as well as its RWD-domain, is required to restrain excessive activation of the eIF2α kinase GCN2. Accordingly, loss of RNF25 results in hyperactivation of GCN2, exacerbated translation shutdown, and impaired cell proliferation following RNA damage—phenotypes that can be fully reversed by genetic or pharmacological inhibition of GCN2. Together, these findings uncover a previously unappreciated RNF25–GCN2 signaling axis and identify ISR-driven toxicity as a potential vulnerability in combination with RNA-damaging chemotherapeutics.

## Introduction

Protein synthesis is a fundamental process required for cellular growth and homeostasis. Because mRNA translation is energetically costly and protein synthesis is prone to errors that can cause toxicity, its global regulation is a central component of cellular adaptation to endogenous and exogenous stress.^1,2^ Indeed, diverse challenges including nutrient deprivation^3^, ultra-violet (UV) radiation^4,5^, metalloid exposure^6^, organelle dysfunction^7,8^, and genotoxic stress^9^, rapidly reprogram global protein synthesis in mammalian cells.

One of the most prominent signaling pathways linking cellular stress to translation repression is the integrated stress response (ISR).^2^ The ISR is activated by four kinases— HRI, PERK, PKR, and GCN2—that converge on phosphorylation of translation initiation factor eIF2α. This modification reduces global protein synthesis by limiting formation of the ternary tRNA^Met^ initiator complex, while simultaneously promoting non-canonical translation of a small set of stress-adaptive mRNAs, including the transcription factor ATF4.^3^ In this way, the ISR enables cells to conserve resources and restore homeostasis. However, excessive or prolonged ISR activation can instead drive growth arrest or cell death, making the magnitude and duration of ISR signaling a key determinant of cell fate.^10^ Although the upstream kinase responsible for ISR activation in response to many stressors has been defined, it has become increasingly clear that context-specific modulation and cross-talk with other pathways are essential to generate appropriately tuned ISR outputs.^5,7,11,12^

Among the four ISR kinases, GCN2 is the evolutionarily most conserved, with budding yeast encoding only a single ISR kinase homolog, Gcn2. Uniquely, GCN2 monitors the translation process itself and is activated in response to a broad range of perturbations including amino acid deprivation, mRNA damage, DNA damage, elongation defects and tRNA mutations.^3,5,9,13,14^ While GCN2 has long been thought to be activated primarily by deacylated tRNAs binding its histidyl-tRNA synthetase–like (HisRS) domain, recent work has suggested that loss of ribosome processivity, resulting in stalling and collision, provides a unifying upstream signal for GCN2 activation.^14–17^ In this framework, the ribosome-associated protein GCN1 serves as a key adaptor that engages GCN2 through a conserved RWD domain and couples ribosome stalling and collision to kinase activation.^16,18^ Despite these recent advances, how such mechanistically distinct perturbations of translation are sensed, integrated, and differentially regulated by the GCN1–GCN2 axis to determine cell fate remains poorly understood. Furthermore, stalled and collided ribosomes themselves pose a major challenge for the cell and are substrates for multiple ubiquitin-mediated quality control pathways in parallel to GCN2 activation, yet how these pathways interface with the ISR to shape appropriate cellular outcomes is also largely unknown.^19^

Here, we use comparative genetic screening across multiple mechanistically distinct perturbations to map the regulatory architecture of acute translation control. These analyzes identify RNF25, a ribosome-associated E3 ubiquitin ligase, as a stress-selective regulator required to sustain protein synthesis following UV irradiation and other RNA-damaging insults. We show that RNF25, in this context, acts independently of its previously described partner RNF14 to restrain excessive GCN2 activation in response to RNA-damage-induced ribosome stalling. Our findings identify RNF25 as a previously unrecognized context-specific suppressor of GCN2 signaling and reveal an antagonistic genetic circuit that preserves translational capacity and cell fitness in response to RNA damage.

## Results

### Systematic profiling of protein synthesis regulators in stress responses

Pulse-labeling cells with the tyrosyl-tRNA mimetic puromycin followed by specific anti-puromycin staining can be used to assess single cell protein synthesis by flow cytometry ^20^ . We recently applied this assay to fluorescent-activated cell sorting (FACS)-based genetic screening to identify SLFN11-driven GCN1/GCN2 activation as a critical node in DNA damage mediated control of protein synthesis.^9^ To further extend these surveys, we applied this approach to a panel of mechanistically distinct stressors - including UV radiation, the endoplasmic reticulum (ER) stressor thapsigargin, and metalloid stress induced by sodium arsenite (Fig. 1A). By screening at intermediate doses and early timepoints (1–2 hours), when translation is attenuated but not fully suppressed (Figs. S1A and S1B), we simultaneously captured both genes whose loss either exacerbated or mitigated stress-induced translation repression, corresponding to positive (blue) and negative (yellow) regulators, respectively (Figs. 1B-1D; Figs. S1C and S1D; Table S1).

**Fig. 1.**
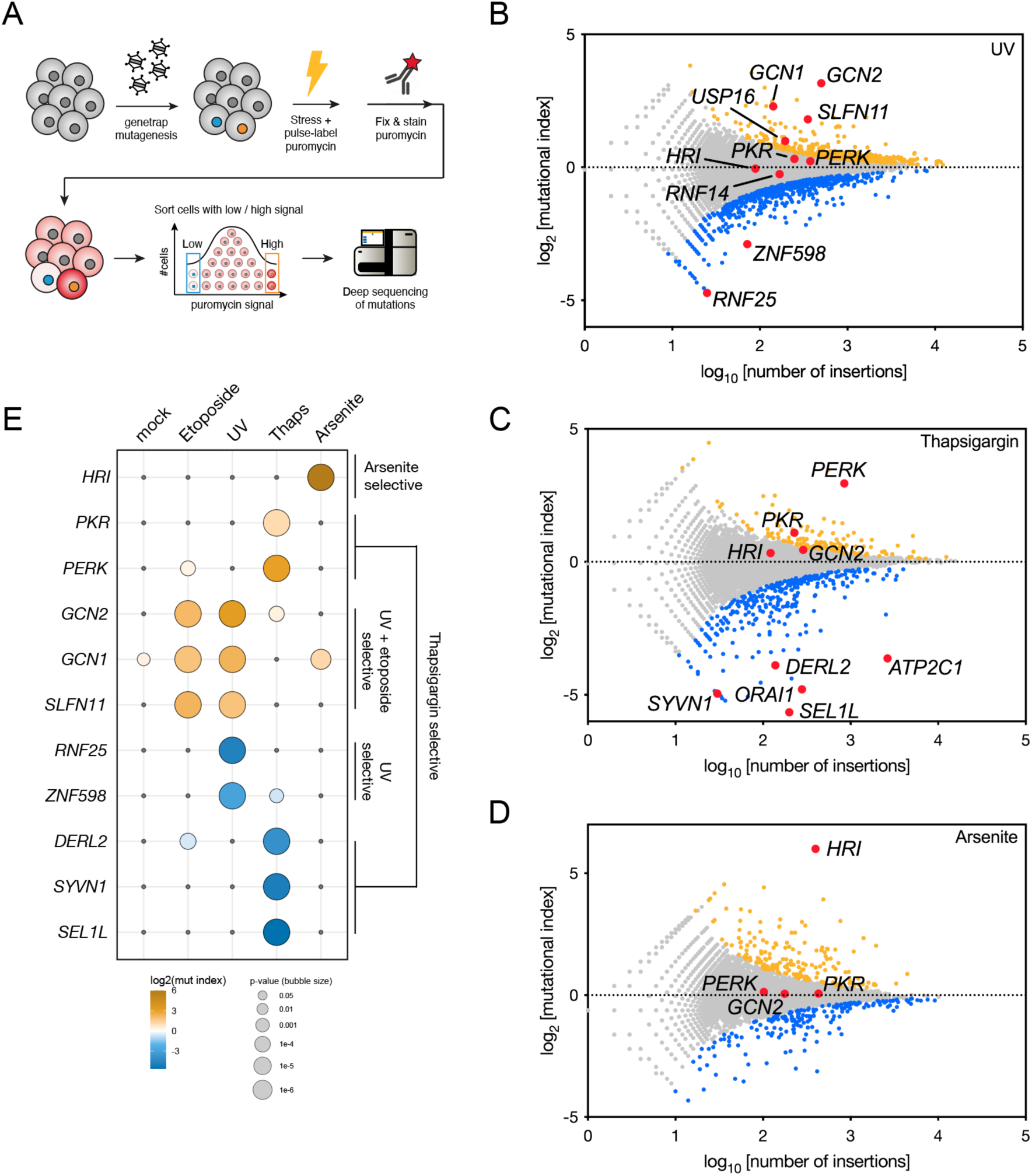
Genome-wide perturbation screens for protein synthesis regulation. (**A**) Workffow schematic of haploid genetic perturbation screens to identify regulators of puromycin incorporation (protein synthesis). (**B-D**) Genetic puromycin incorporation screens after treatment with (B) UV (25J/m^2^, 2 hr recovery), (C) thapsigargin (4 μM, 1.5 hr) or (D) sodium arsenite (100 μM, 1 hr), presented as fishtail plots. Prior to harvesting, cells were pulse-labeled with puromycin (2 μg/ml) for 15 min. Genes are plotted according to their mutational index (MI, y axis) and the total number of gene-trap insertions (x axis), with significant positive and negative regulators colored in blue and yellow, respectively [two-sided Fisher’s exact test, false discovery rate (FDR) corrected P-value ≤ 0.05; non- significant genes are shown in grey]. Selected genes of interest for each screen are highlighted in red for visual purposes. See Table S1 for full list of regulators. (**E**) Bubble heat map of selected genes to illustrate comparative identification of perturbation- specific regulators. Genes are plotted according to their MI (color) and significance (size) across five independent puromycin incorporation screens (Figs. 1B-D and Figs. S1C and S1D).^9^

Across conditions, these comparative screens recovered core components of translational control and the ISR, including all four ISR kinases (GCN2, PERK, HRI, and PKR) as significant regulators, validating the robustness of the approach (Fig. 1E). Importantly, the screens also revealed stress-specific regulatory signatures among a wealth of potential new regulators. For example, in line with thapsigargin’s inhibitory effect on Ca^2+^ sequestration in the ER, the screen selectively identified the ER-resident ISR kinase PERK as a negative regulator, along with genes involved in ER calcium homeostasis (e.g. *ATP2C1* and *ORAI1*) and ER-associated degradation (ERAD, e.g. *SEL1L, SYVN1* and *DERL2*) as positive regulators, consistent with established ER stress biology (Fig. 1C and 1E; Table S1).^8,21^ Thus, these data demonstrate that acute translational control is wired in a highly context-dependent manner that can be uncovered by comparative puromycin incorporation screens.

Notably, while the UV screen shared many regulators with a previous puromycin screen for DNA damage induced by etoposide - including the SLFN11-GCN1/GCN2 pathway - it also revealed a number of UV-selective regulators of protein synthesis, most prominently *RNF25* as the top positive UV regulator (Figs. 1B and 1E; Fig. S1D).^9^ Furthermore, underlining *RNF25’s* specificity for this phenotype, *RNF25* did not score as a significant regulator in more than 30 publicly available HAP1 phenotypic screens covering various biological processes (Fig. S1E). RNF25 is linked to the translational machinery as a reported ribosome associated ubiquitin ligase, however, how RNF25 promotes global protein synthesis is not known.^22,23^ We therefore focused our efforts on RNF25 to characterize its role in sustaining translation in the UV-induced stress response.

### RNF25 ubiquitin ligase activity promotes translation in response to UV irradiation independently of RNF14

Prompted by our genetic screening results, we generated HAP1 cell lines with targeted deletion of RNF25 (Δ*RNF25*) to investigate how RNF25 regulates protein synthesis following UV irradiation. In wild-type HAP1 cells, 25 J/m^2^ UV induced an intermediate translational shutdown two hours post irradiation, characterized by a bimodal distribution of cells with high and low puromycin incorporation, as assessed by flow cytometry (Fig. 2A; Fig. S1B). This bimodal on/off response is characteristic of the SLFN11-driven pathway activated by DNA damage, which exhibits threshold-like activation that is strongly influenced by cell state, including cell-cycle position.^9,24^ While loss of RNF25 did not measurably affect nascent protein synthesis under mock-treated conditions, UV-treated Δ*RNF25* cells exhibited a markedly enhanced translation shutdown compared to wild-type cells, validating our genetic screening results (Fig. 2A; Fig. S2A). Furthermore, time-resolved analysis of bulk protein synthesis by immunoblot showed that the exacerbated UV-induced decrease of translation in Δ*RNF25* cells was detectable as early as 15 min post irradiation indicating that RNF25 functions in the acute phase of the UV response (Fig. 2B).

**Fig. 2.**
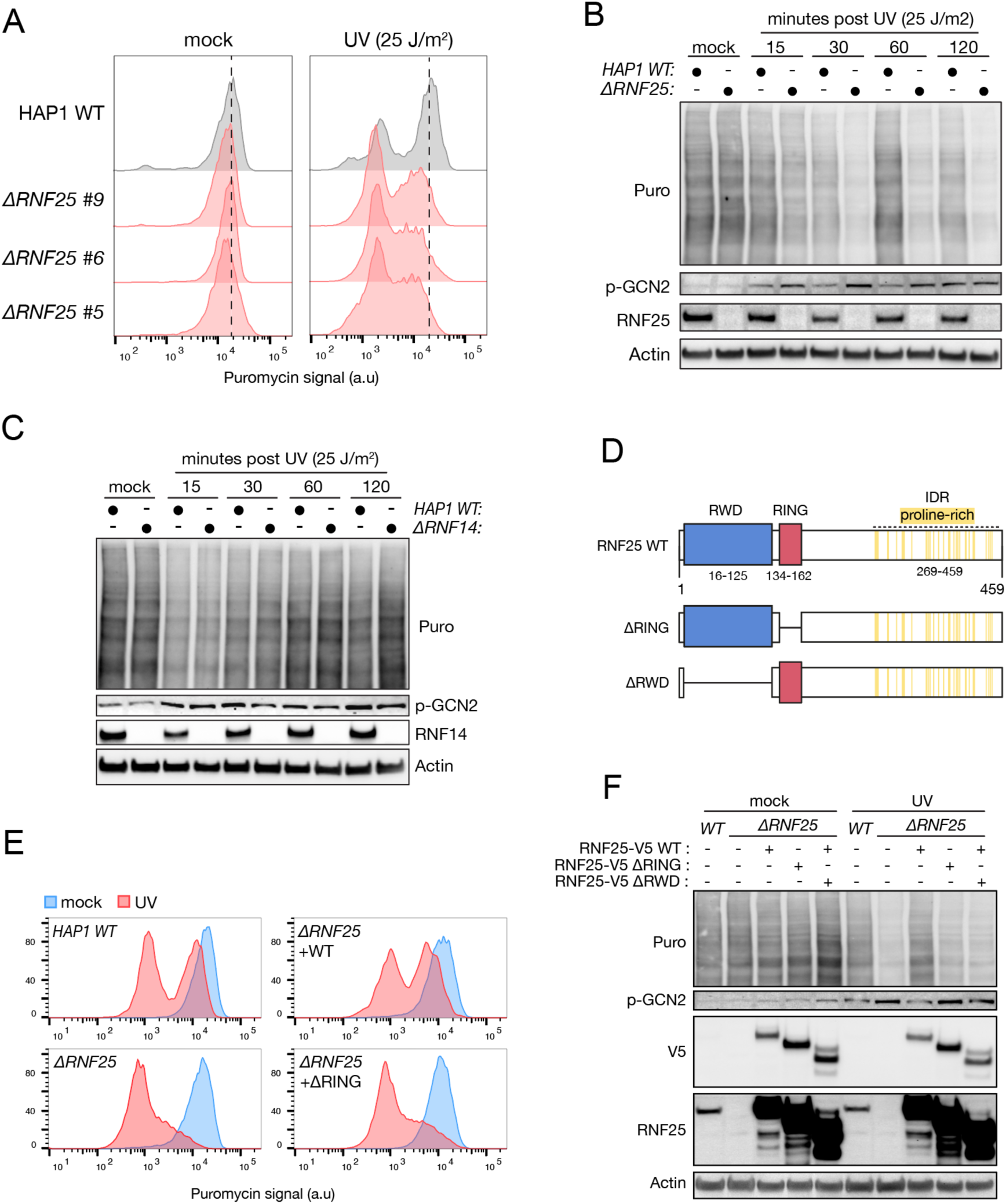
RNF25 ubiquitin ligase activity promotes translation in response to UV radiation independently of RNF14. (**A)** Flow cytometry analysis of puromycin incorporation after mock or UV treatment (25J/m^2^, 2 hr recovery) in parental HAP1 and three independent RNF25 knockout (ΔRNF25) clones. See also Fig. S2A. (**B**) As in (A) but cells subjected to immunoblot analysis to monitor bulk puromycin incorporation at indicated timepoints following UV treatment. (**C**) As in (B) but comparing parental HAP1 wild-type (WT) and RNF14 knock- out (ΔRNF14) cells. (**D**) Schematic of human RNF25 wild-type (WT) domains and deletion mutants. (**E-F**) Treated as in (A) but comparing parental HAP1, ΔRNF25 and ΔRNF25 cells complemented with ectopic RNF25-V5 wild-type (WT), RING domain deletion mutant (ΔRING) or RWD-domain deletion mutant (ΔRWD). Cells were subjected to (E) flow cytometry analysis or (F) immunoblot analysis using the indicated antibodies.

RNF25 has recently been linked to an RNF14-dependent ubiquitination pathway in ribosome surveillance, however, RNF14 did not score as a significant regulator in our genetic UV screen (Fig. 1B).^22,23,25^ Consistent with this, targeted deletion of *RNF14* in HAP1 cells (Δ*RNF14*) did also not affect global protein synthesis following UV treatment compared to wild-type cells (Figs. 2C; Fig. S2B). These data indicate that RNF25 operates independently of the RNF25-RNF14 pathway to regulate UV-induced translation control.

RNF25 is also an E3 ubiquitin ligase and contains an N-terminal RWD domain, similar to GCN2, and a RING domain required for substrate ubiquitination (Fig. 2D).^25,26^ To test whether these domains were required for its new function in translation control, we utilized a lentiviral complementation system allowing us to reintroduce ectopic V5- tagged RNF25 constructs into Δ*RNF25* HAP1 cells. Importantly, Δ*RNF25* cells complemented with RNF25-V5 wild-type (WT) restored translation repression to levels comparable to parental HAP1 cells, whereas deletion of the RWD- (ΔRWD, amino acids 16-125) or RING-domain (ΔRING, amino acids 134-165) abolished this rescue despite similar expression (Figs. 2E and 2F).

We conclude that RNF25 requires its RWD-domain and ubiquitin ligase activity, but not RNF14, to sustain protein synthesis following UV irradiation.

### RNF25 and SLFN11 modulate translation in parallel following UV radiation

Both our genetic screen(s) and previous work have implicated SLFN11-induced ribosome stalling and GCN1-GCN2 signaling in the bimodal translation shutdown following DNA damage, such as that induced by etoposide and UV radiation (Figs. 1B and 1E; Fig. S1D).^9^ However, RNF25 scored as a positive regulator of translation exclusively in our UV screen and not in the etoposide screen, arguing against a general role for RNF25 in suppressing DNA damage– or SLFN11-mediated ribosomal stalling (Fig. 1E).

To directly test whether the enhanced UV-induced translation shutdown observed in Δ*RNF25* cells depends on SLFN11, we generated HAP1 cells lacking SLFN11 (Δ*SLFN11*), as well as double knockout cells (Δ*SLFN11* Δ*RNF25*). As expected, loss of SLFN11 largely abolished the bimodal translation shutdown induced by both UV and etoposide treatment (Figs. 3A and 3B; Figs. S3A and S3B). In contrast, loss of RNF25 selectively impaired protein synthesis in response to UV but not etoposide (Figs. 3A and 3B; Figs. S3A and S3B). Importantly, Δ*SLFN11* Δ*RNF25* cells still underwent a pronounced translation shutdown following UV irradiation, but not following etoposide treatment, demonstrating that RNF25 regulates a SLFN11-independent pathway controlling protein synthesis in response to UV radiation (Figs. 3A and 3B; Figs. S3A and S3B).

**Fig. 3.**
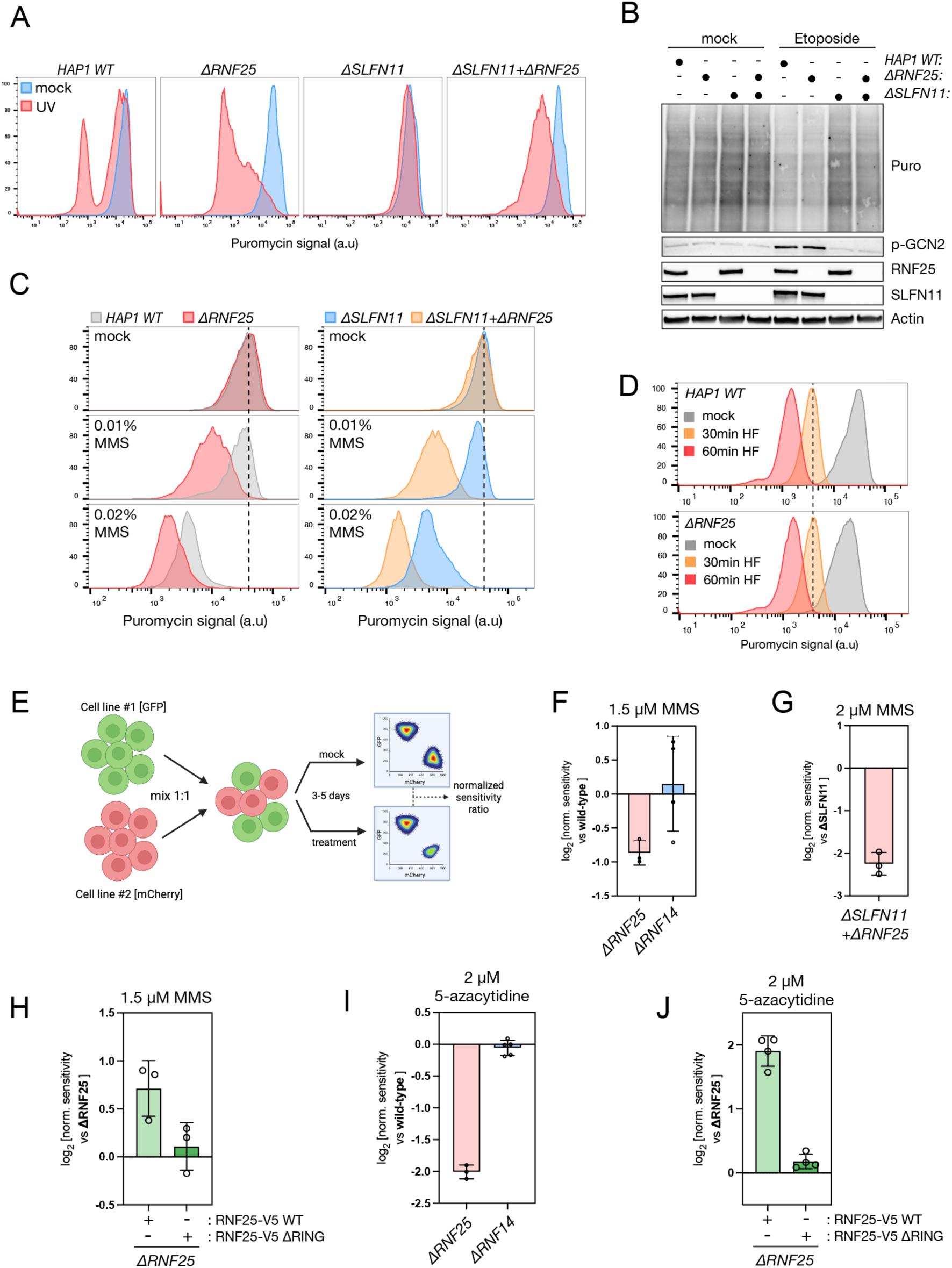
RNF25 is required for protein synthesis and cell proliferation in response to RNA damage. (**A**) Flow cytometry analysis of puromycin incorporation after mock or UV treatment (25J/m^2^, 2 hr recovery) in parental HAP1 cells and genetic knock-out cell lines (Δ) for RNF25, SLFN11 and RNF25+SLFN11. (**B**) As in (A) but cells were treated with etoposide (5 μM, 3 hrs) and subjected to immunoblot analysis to monitor bulk puromycin incorporation. (**C**) As in (A) but cells were treated with indicated concentrations of MMS for 1hr. (**D**) As in (A) but cells were treated with 1 μM halofugione for 30 and C0 mins. (**E**) Schematic of growth competition assay using ffuorescently labeled cell lines. (**F**) Growth competition assay comparing proliferation of parental HAP1 cells to ΔRNF25 and ΔRNF14 cells, respectively, in mock and MMS-treated conditions (1.5 μM) for three days. Data represented as mean±SD of the normalized drug sensitivity ratio (log2). n = 3-4 independent biological replicates. (**G**) As in (F) but comparing HAP1 ΔSLFN11cells to ΔSLFN11ΔRNF25 cells treated with 2 μM MMS for five days. Data represents mean±SD, n = 4 independent biological replicates. (**H**) As in (F) but comparing ΔRNF25 cells to ΔRNF25 cells complemented with ectopic RNF25 wild-type (WT) or a RING deletion mutant (ΔRING), respectively. Data represents mean±SD, n = 3 independent biological replicates. (**I**) As in (F) but treating cells with 2 μM 5-Azacytidine for three days. Data represents mean±SD, n = 3-4 independent biological replicates. (**J**) As in (H) but cells were treated with 2 μM 5-Azacytidine for three days. Data represents mean±SD, n = 4 independent biological replicates.

Notably, the translation shutdown observed in Δ*SLFN11* Δ*RNF25* cells treated with 25 J/m^2^ UV lacked the bimodal puromycin incorporation profile characteristic of SLFN11- driven regulation and instead manifested as a homogenous unimodal decrease across the entire cell population (Fig. 3A). A similar unimodal puromycin decrease was observed in Δ*SLFN11* cells treated with higher doses of UV irradiation, confirming SLFN11- independent regulation in the UV response also in RNF25-proficient cells (Fig. S3C). These observations further support the notion that RNF25 governs a mechanistically distinct origin or mode of translation control compared to DNA damage activated SLFN11.

Because many commonly used cancer and immortalized cell lines lack SLFN11 expression, and thus, a functional pathway connecting DNA damage to ribosome stalling, we hypothesized that such cell lines would display a unimodal, RNF25- dependent translational response to UV.^27,28^ Consistent with this prediction, SLFN11- negative wild-type HEK293T cells exhibited a unimodal translation shutdown at higher UV doses, which was further exacerbated by RNF25 ablation (Figs. S3D and S3E).

Together, these results demonstrate that RNF25 is required to sustain protein synthesis following UV irradiation through a mechanism that operates in parallel to, but independently of, SLFN11-mediated translation repression triggered by DNA damage.

### RNF25 promotes translation in response to RNA damaging conditions

Given the selective requirement for RNF25 in sustaining translation following UV irradiation, but not etoposide treatment, we next asked which UV-specific lesions underlie this effect. We reasoned that this difference could reflect UV-induced crosslinking of RNA, in contrast to etoposide, which primarily induces DNA double-strand breaks through highly specific inhibition of DNA topoisomerase II cleavage complexes.^29,30^ Indeed, recent work has demonstrated that UV-induced lesions in mRNA templates can induce ribosome stalling and that many aspects of the acute cellular response to UV irradiation are driven by RNA- rather than DNA-damage.^31–33^ Based on this distinction, we hypothesized that RNF25 promotes protein synthesis by regulating translational responses to RNA damage, but not other stimuli inducing ribosomal stalling. To identify additional RNA damaging conditions that rely on this new RNF25 pathway we employed the alkylating agent methyl methanesulfonate (MMS) that also damages both DNA and RNA molecules and is known to cause ribosome stalling.^14,29^ Indeed, MMS treatment caused an acute dose-dependent shutdown of translation which was further exacerbated by loss of RNF25 (Fig. 3C; Figs. S4A and S4B). Interestingly, the puromycin incorporation profile, as assessed by flow cytometry, was unimodal and SLFN11 loss had minimal effect on the translation shutdown at the tested timepoints and concentrations of MMS (Fig. 3C; Fig. S4B). Thus, these data also indicate that, at least during the acute phase, MMS preferentially impacts on RNF25-dependent translation regulation rather than activating the SLFN11-driven pathway, in contrast to UV irradiation.

Next, we sought a condition that induces ribosomal stalling and translation shutdown independently of RNA or DNA damage. To this end, we used halofugione, a prolyl-tRNA synthetase inhibitor that causes rapid accumulation of uncharged tRNA^Pro^, ribosome stalling and collision, and GCN2-dependent translational repression.^13,14^ Consistent with the RNA damage hypothesis, wild-type and Δ*RNF25* HAP1 cells exhibited a comparable, time-dependent repression of protein synthesis following halofugione treatment (Fig. 3D).

Together, these findings indicate that RNF25 is selectively required for sustaining translation in response to RNA damaging conditions.

### RNF25 promotes cell proliferation in response to RNA damaging conditions

Next, we asked how loss of RNF25 would impact the cellular response to RNA damage. To this end, we used an internally controlled growth competition assay by fluorescently labeling HAP1 cell lines and assessing their relative proliferation in co-culture experiments by flow cytometry (Fig. 3E).^34^ We initially focused on MMS because it elicited a prominent RNF25-dependent translation phenotype with minimal SLFN11 contribution in the acute response (Fig. 3C). Interestingly, cells deleted for RNF25 proliferated slower than wild-type HAP1 cells in response to MMS treatment (Fig. 3F). Notably, this growth defect became even more pronounced when we genetically eliminated the SLFN11-dependent death pathway which allowed us to increase the MMS dose also indicating that DNA damage and SLFN11-driven responses are also induced by long term MMS treatments (Fig. 3G). These data are in line with previous CRISPR-Cas9 genetic screens linking loss of RNF25 to MMS sensitivity in human cells.^34,35^ Moreover, aligning with the genetics of our molecular observations, we confirmed that MMS sensitivity was independent of RNF14 status and that loss of RNF25 did not give rise to etoposide sensitivity at a dose where Δ*SLFN11* cells were markedly resistant (Figs. 3F and S4C). Importantly, by comparing Δ*RNF25* cells directly to Δ*RNF25* cells complemented with ectopic RNF25, we confirmed the requirement for the RNF25 RING domain in supporting cell proliferation across MMS-treated conditions (Fig. 3H).

Prompted by these findings, we looked for more clinically relevant drugs which could modify or damage RNA more selectively to assess if RNF25 status could be relevant in anti-cancer treatment. The widely used chemotherapeutic anti-metabolites 5- Fluorouracil (5-FU) and 5-Azacytidine have been shown to incorporate mainly into RNA over DNA, but the relevance of this for anti-cancer treatment is still not well understood.^36–38^ Although we did not observe any robust sensitivity of Δ*RNF25* cells toward 5-FU (data not shown), we noticed a striking proliferation defect after 5- Azacytidine treatment compared to wild-type cells (Fig. 3I). Importantly, this growth defect required the RNF25 RING domain and was independent of RNF14, suggesting a mechanism analogous to that of MMS (Figs. 3I and 3J).

In summary, the here tested RNA damaging stressors challenge translation control and cell proliferation by similar mechanisms that requires RNF25 ubiquitin ligase function.

### RNF25 promotes translation and cell proliferation by curbing GCN2 hyperactivation

In both the UV and MMS, but not etoposide, experiments we noticed a robust RNF25- dependent increase in GCN2 activation already at the early timepoints as monitored by phospho-specific immunoblotting of the well-established GCN2 Thr899 autophosphorylation mark (Figs. 2B, 2F and 3B; Figs. S2A, S3A, S4A and S4B). Importantly, this increase was dependent on the RING- and RWD-domain of RNF25 but not observed in UV-treated RNF14 deficient cells uncoupling this phenotype from the RNF25-RNF14 pathway (Figs. 2C and 2F). This prompted us to ask the question if acute hyperactivation of GCN2 and the ISR could account for the enhanced translation shutdown observed in RNF25 deficient cells. To test this, we generated cells with targeted deletion of GCN2 in HAP1 wild-type- (Δ*GCN2*) and Δ*RNF25*-background (Δ*GCN2* Δ*RNF25*). Strikingly, GCN2 loss completely abrogated the enhanced UV-induced translation shutdown in RNF25 deficient cells (Fig. 4A; Fig. S5A). To corroborate these findings, we treated cells using chemical inhibitors of GCN2 (GCN2i, A-92) and the ISR (ISRIB) which phenocopied the loss of GCN2 by abolishing the translation shutdown in Δ*RNF25* cells (Fig. 4B; Fig. S5B). We conclude that increased GCN2 signaling and ISR activation is responsible for the enhanced translation shutdown observed in RNF25 deficient cells.

**Fig. 4.**
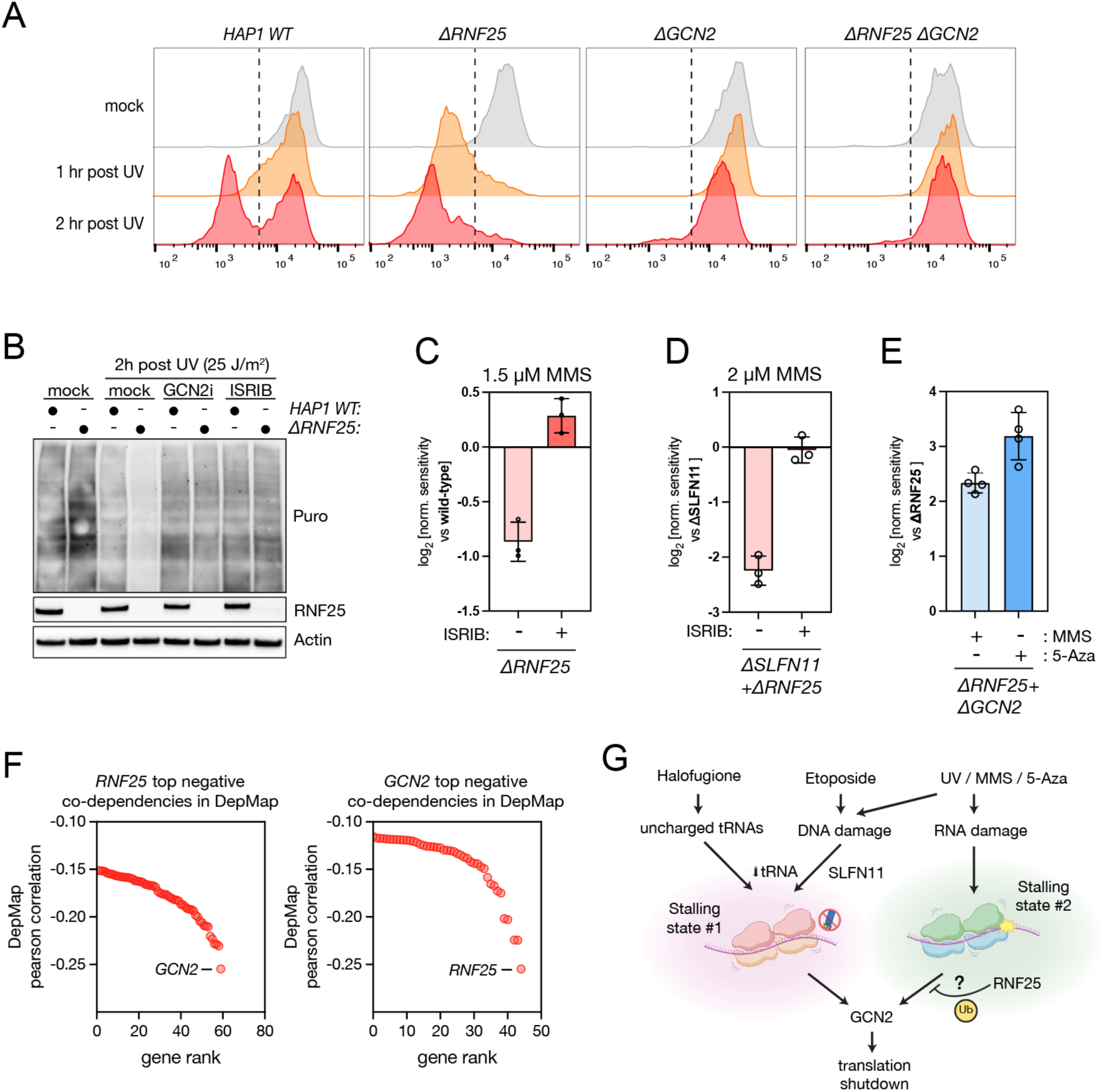
RNF25 suppresses toxic GCN2 hyperactivation in response to RNA damage. (**A**) Time-resolved ffow cytometry analysis of puromycin incorporation after mock and UV treatment (25J/m^2^) in parental HAP1 cells and genetic knock-out (Δ) cells for RNF25, GCN2 and RNF25+GCN2. (**B**) Parental HAP1 and ΔRNF25 cells were mock or UV-treated (25J/m^2^, 2hr recovery) in the presence or absence GCN2 inhibitor (GCN2i A-S2, 1 μM) or the ISR inhibitor ISRIB (0.2 μM), respectively. Cells were then labeled with puromycin and subjected to immunoblot analysis using the indicated antibodies. (**C**) Growth competition assay comparing proliferation of parental HAP1 cells to ΔRNF25 in mock and MMS-treated conditions (1.5 μM) in the presence of ISRIB (0.2 μM) or not, for three days. Data represented as mean±SD of the normalized drug sensitivity ratio (log2). n = 3 independent biological replicates. Samples not treated with ISRIB are also plotted in Fig. 3F. (**D**) As in (F) but comparing HAP1 ΔSLFN11 cells to ΔSLFN11ΔRNF25 cells treated with 2 μM MMS for five days. Data represents mean±SD, n = 4 independent biological replicates. Samples not treated with ISRIB are also plotted in Fig. 3G. (**E**) As in (C) but comparing ΔGCN2 cells to ΔGCN2+ΔRNF25 cells cultured in media containing 1 μM MMS or 2 μM 5-Azacytidine (5-Aza) as indicated. Data represents mean±SD, n = 4 independent biological replicates. (**F**) Co-dependency data exported from Cancer Dependency Map (CRISPR DepMap Public 24Q4). The exported top negative co-dependencies for RNF25 (left) and GCN2 (right) were plotted and ranked according to their DepMap pearson correlation score. (**G**) Model of RNF25 selectivity

Considering these observations, we next asked if hyperactivation of GCN2 and ISR is a protective or maladaptive response for cell proliferation. Remarkably, co-treating cells with the ISR inhibitor ISRIB completely abolished the proliferation defect observed in MMS-treated Δ*RNF25* cells, independently of SLFN11 status (Figs. 4C and 4D). These data were further supported by the observation that GCN2 loss in Δ*RNF25* cells lead to a pronounced resistance to MMS- and 5-Azacytidine treatments (Fig. 4E). Collectively, these data indicate that the principal function of RNF25 in supporting cell fitness upon RNA damage is to curtail toxic GCN2 hyperactivation.

Based on recent work showing that ribosome stalling and collisions are necessary but not sufficient for GCN2 activation via GCN1, we considered two non-mutually exclusive scenarios for how RNF25 loss impacts on GCN2 signaling.^14,17^ First, RNF25 loss may increase ribosome stalling and collision frequency, for example by impairing collision resolution or the repair of RNA lesions, as have been shown to enhance GCN2 signaling in yeast.^39^ Alternatively, RNF25 could directly modulate GCN2 signaling downstream of, or in conjunction with, stalled or collided ribosomes. To distinguish between these possibilities, we performed disome analysis by polysome profiling, which revealed evident UV-induced disome formation 30 minutes after irradiation (Fig. S6A). However, loss of RNF25 did not measurably increase disome abundance, despite pronounced GCN2 hyperactivation and translational shutdown already at this time point (Figs. 2B; Figs. S3A and S6A).

Consistent with ribosome collision accumulation not being sufficient to drive GCN2 hyperactivation in our system, ZNF598, a gene that scored less prominently in our UV screen than RNF25, is a well-established sensor of collided ribosomes that promotes their resolution (Fig. 1B).^40–43^ Indeed, although Δ*ZNF598* cells did display a modest increase in translational repression following UV-treatment, as suggested by our screen, they did not exhibit elevated GCN2 activation (Figs. S6B and S6C), indicating that impaired collision resolution alone does not phenocopy RNF25 loss. Together, these data argue that excessive accumulation of ribosome collisions is not the primary driver of GCN2 hyperactivation in RNF25-deficient cells.

The observation that both GCN2 deletion and pharmacological ISR inhibition completely suppress RNF25-dependent phenotypes further supports the idea that RNF25 functions within the GCN2 signaling pathway acting downstream of, or in parallel to, RNA damage and ribosomal stalling. Remarkably, the strong genetic interaction observed in our study is independently supported by the Cancer Dependency Map where *RNF25* and *GCN2* emerge as each other’s top negative co-dependencies in CRISPR-Cas9 fitness screens across more than a thousand human cell lines originating from diverse tissues and cancers (Fig. 4F).^44^ This pattern is characteristic of genes that function in antagonistic pathways, consistent with RNF25 acting to restrain GCN2 activity. Furthermore, this suggests that the molecular function and relationship between RNF25 and GCN2 is not only important in conditions of artificially induced RNA damage stress but also relevant across unperturbed proliferation of human cancer cell lines.

## Discussion

In this study, we define a previously unrecognized RNF25-GCN2 signaling axis that sustains protein synthesis and promotes cell proliferation in response to RNA damage. Using comparative genetic screening and mechanistic dissection, we identify RNF25 as a stress-selective regulator that limits excessive activation of the ISR kinase GCN2. Our data support a model in which RNF25 does not broadly suppress GCN2 signaling induced by stalled ribosomes—such as those generated by uncharged or SLFN11-cleaved tRNAs—but instead selectively restrains GCN2 in response to ribosomes stalled by RNA damage (Fig. 4G). The basis for this selectivity remains to be determined but may involve structurally distinct stalled/collided ribosome states - e.g., differences in A-site occupancy, decoding geometry, or collision architecture - that differentially engage RNF25 and/or GCN2. Consistent with this idea, the underlying RNA lesions are likely different across our conditions: UV induces bulky photolesions and RNA–protein crosslinks, whereas MMS predominantly generates alkylated RNA bases, yet both can stall elongating ribosomes on damaged mRNA templates.^29^

How might RNF25 mechanistically restrain GCN2 activation? Two previously described RNF25 functions coupled to ribosome surveillance – degradation of RNA–protein crosslinks and proteins trapped in the ribosomal A-site - require RNF14^22,23,25^ ; however, our data clearly show that RNF14 is dispensable for RNF25-dependent regulation of GCN2 (Fig. 2C, 3F, and 3I; Fig. S2B). These data demonstrate that RNF25 has other independent functions in addition to supporting the RNF25-RNF14 pathway in ribosome surveillance. GCN2 activation requires engagement of GCN1 via a conserved RWD domain and other RWD-containing proteins, such as the reported GCN2 suppressor IMPACT, have been proposed to compete for GCN1 binding and thereby inhibit GCN2 activation.^45,46^ Because RNF25 also contains a conserved RWD domain required for translation control (Fig. 2D), one possibility is that loss of RNF25 could simply promote GCN1-GCN2 complex formation by lowering the competition for GCN1-binding. However, neither IMPACT, DFRP2, or RNF14 scored as regulators of protein synthesis in the UV screen although they all have validated GCN1-binding RWD domains and are expressed at higher levels than RNF25 in HAP1 cells (Fig. 1B; Fig. S7A; Table S1).^25,45,46^ Moreover, expressing ectopic RNF25 WT manyfold over endogenous levels did not further suppress GCN2 activation or inhibit translation repression, arguing against a simple competitive GCN1-binding model (Figs. 2F). Such a competition model would also not explain the requirement for RNF25’s RING domain in translation control, an observation that rather implicates ubiquitin signaling in this process. Another non-mutually exclusive possibility is therefore that RNF25’s RWD domain facilitates transient recruitment to GCN1-bound ribosomes stalled or collided by RNA damage which enable RNF25-mediated ubiquitylation of ribosome-associated components or regulatory factors in a manner that constrains GCN2 activation efficiency. Consistent with this idea, full GCN2 activation, in addition to GCN1 binding, relies on multiple specific interactions with the ribosome, which could be regulated by site-specific ubiquitin modifications.^14,47–50^ We therefore propose that RNF25 regulates GCN2 signaling through direct ubiquitination of ribosomal substrates, with RPS27A Lys113 emerging as a particularly compelling candidate. This site is mono-ubiquitinated in an RNF25-dependent manner and is required for activation of RNF14, raising the possibility that the same modification may also tune GCN2 activation on the ribosome in certain context.^25^ Supporting this model, USP16—the deubiquitinating enzyme that removes mono-ubiquitin from RPS27A Lys113—scored as a negative regulator of protein synthesis in our UV screen, the opposite effect of RNF25 loss (Fig. 1B).^51^ Together, these observations point to RPS27A Lys113 ubiquitination as a potential regulatory node linking RNF25 activity to GCN2 control. Future work will be required to define the full scope of relevant ubiquitin substrates and determine how these modifications structurally intersect with GCN2 activation on the ribosome.

Beyond its mechanistic implications, our work has potential clinical relevance. The ribonucleoside analogue 5-Azacytidine is widely used as a primary treatment of myelodysplastic syndromes and acute myeloid leukemia.^52^ Although 5-Azacytidine’s mode of action is often framed as a DNA hypomethylating agent, the majority (80-90%) of 5-Azacytidine is actually incorporated into RNA, yet, the importance of this in cancer cell killing is not understood.^36,52^ Our findings suggest that RNF25 protects cells from toxic GCN2 hyperactivation during RNA damage and that disruption of this pathway sensitizes cells to 5-Azacytidine (Figs. 3I, 3J and 4E). In this context, the exact RNA lesion(s) caused by 5-Azacytidine remains to be defined but we propose they would induce ribosome stalling analogous to MMS and UV-induced lesions in mRNA templates. These observations also raise the possibility that inhibition of RNF25 or pharmacological activation of GCN2, for which there are several compounds available, could enhance the efficacy of 5-Azacytidine and potentially other RNA modifying chemotherapeutics.^53^ Supporting the broader notion of RNA damage as an important aspect of anti-cancer treatment, it was recently shown that the cell death response towards 5-FU, one of the most used agents against solid cancers, depend largely on incorporation into ribosomal RNA (rRNA), and not DNA.^37,38^ Collectively, these studies indicate that RNA damage may represent an overlooked aspect of cell killing for several nucleic acid targeting chemotherapeutics which could be further exploited to enhance their anti-cancer effect.

Finally, RNF25 and GCN2 exhibit a clear antagonistic genetic interaction across cancer cell lines in the Cancer Dependency Map (Fig. 4F)^44^, indicating that this regulatory axis is broadly relevant for cellular fitness even in unperturbed growth conditions. In the future it will be important to define the endogenous signals underlying this genetic interaction and study the physiological role of the RNF25-GCN2 signaling axis in cells and organisms.

### Limitations of this study

While RNF25 loss did not measurably increase the bulk ribosome collision burden in our conditions, we cannot exclude qualitative or localized changes in stalled ribosome structures that may influence GCN2 signaling. Our study is also limited to cell-culture models and acute stress responses, and the relevance of this pathway in tissues or *in vivo* remains to be explored. Finally, the specific RNA lesions or ribosome-associated signals that engage the RNF25–GCN2 axis are not yet defined and may not be exclusive to RNA damage.

## Acknowledgements

We thank Thijn Brummelkamp for sharing reagents related to GCN2 and SLFN11 as well as the staff of the SUND Genomics platform (University of Copenhagen), especially Heike Wollmann and Magali Michaut, for support in establishing a Next Generation Sequencing pipeline for HAP1 mutagenesis screens. Work in the Haahr lab was supported by the Lundbeck foundation (R445-2023-1054) and the Kirsten & Freddy Johansens foundation. Center for Gene Expression (CGEN) is a Center of Excellence funded by The National Danish Research Foundation (grant no. DNRF166).

## Resource availability

### Lead contact

Correspondence and requests for materials should be addressed to Peter Haahr.

### Data and code availability

The sequencing data for genetic puromycin incorporation screens have been deposited at the NCBI Sequence Read Archive (PRJNA1415547). All significant positive and negative regulators from each screen are available in Supplementary Table S1. The Bioinformatic pipeline used for screen analysis is available at GitHub (https://github.com/BrummelkampResearch/phenosaurus).

### Author contributions

A.S.S. and P.H. conceived and designed the study. A.S.S., L.N., M.G.-T., X.S., S.R., and K.C.M. performed experiments and analyzed data. P.H. and S.B.-J. supervised experiments. P.H. wrote the manuscript. All authors commented on and edited the manuscript.

## Supplemental Figs

**Fig. S1.**
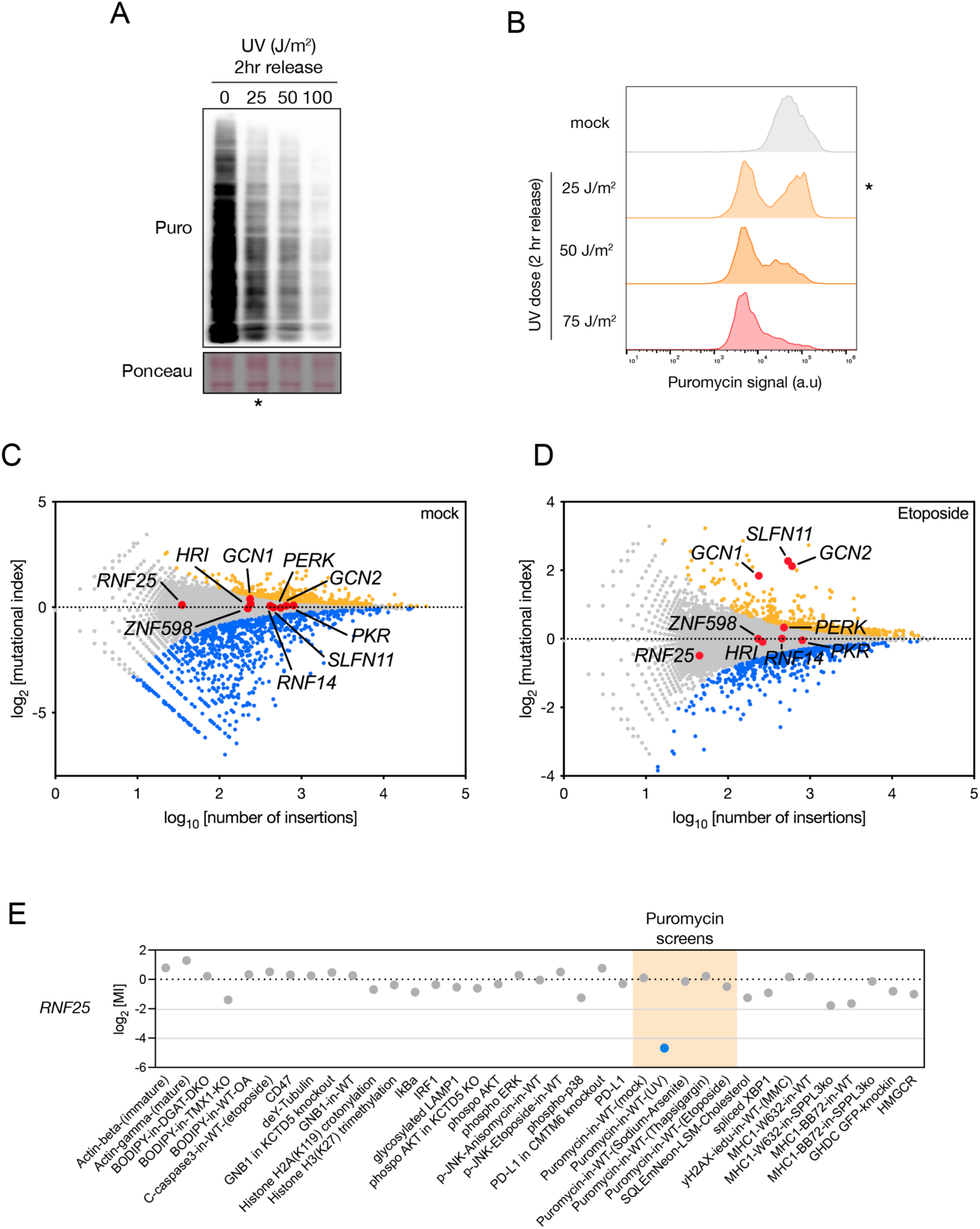
Genome-wide perturbation screens for protein synthesis regulation. (**A-B**) Example of screen perturbation dose titration. Parental HAP1 cells treated with the indicated dose of UV were allowed to recover for two hours before puromycin labeling and analysis by (A) immunoblot and (B) flow cytometry analysis to monitor puromycin incorporation. Asterisk (*) indicates the condition used in the genetic screen (Fig. 1B) (**C-D**) Genetic puromycin incorporation screens after (A) mock treatment or (B) Etoposide treatment (5 μM, 3h), presented as fishtail plots. Prior to harvesting, cells were pulse-labeled with puromycin (2 μg/ml) for 15 min. Genes are plotted according to their mutational index (MI, y axis) and the total number of gene-trap insertions (x axis), with significant positive and negative regulators colored in blue and yellow, respectively [two-sided Fisher’s exact test, false discovery rate (FDR) corrected P-value ≤ 0.05; nonsignificant genes are shown in grey]. Selected genes for each screen are highlighted in red (corresponding to the genes highlighted in Fig. 2B). Data were acquired from Boon et al. 2024.^9^ (**E**) RNF25 mutational index (MI) scores from published HAP1 phenotypic screens. Based on the FDR-corrected P-values ≤ 0.05, non-significant MI scores are colored in grey (36 screens) and significant in blue (1 screen; Puromycin-in-WT-(UV)). RNF25 MI scores and p-values were acquired from published screens as in as in Bianchi et al.^54^

**Fig. S2.**
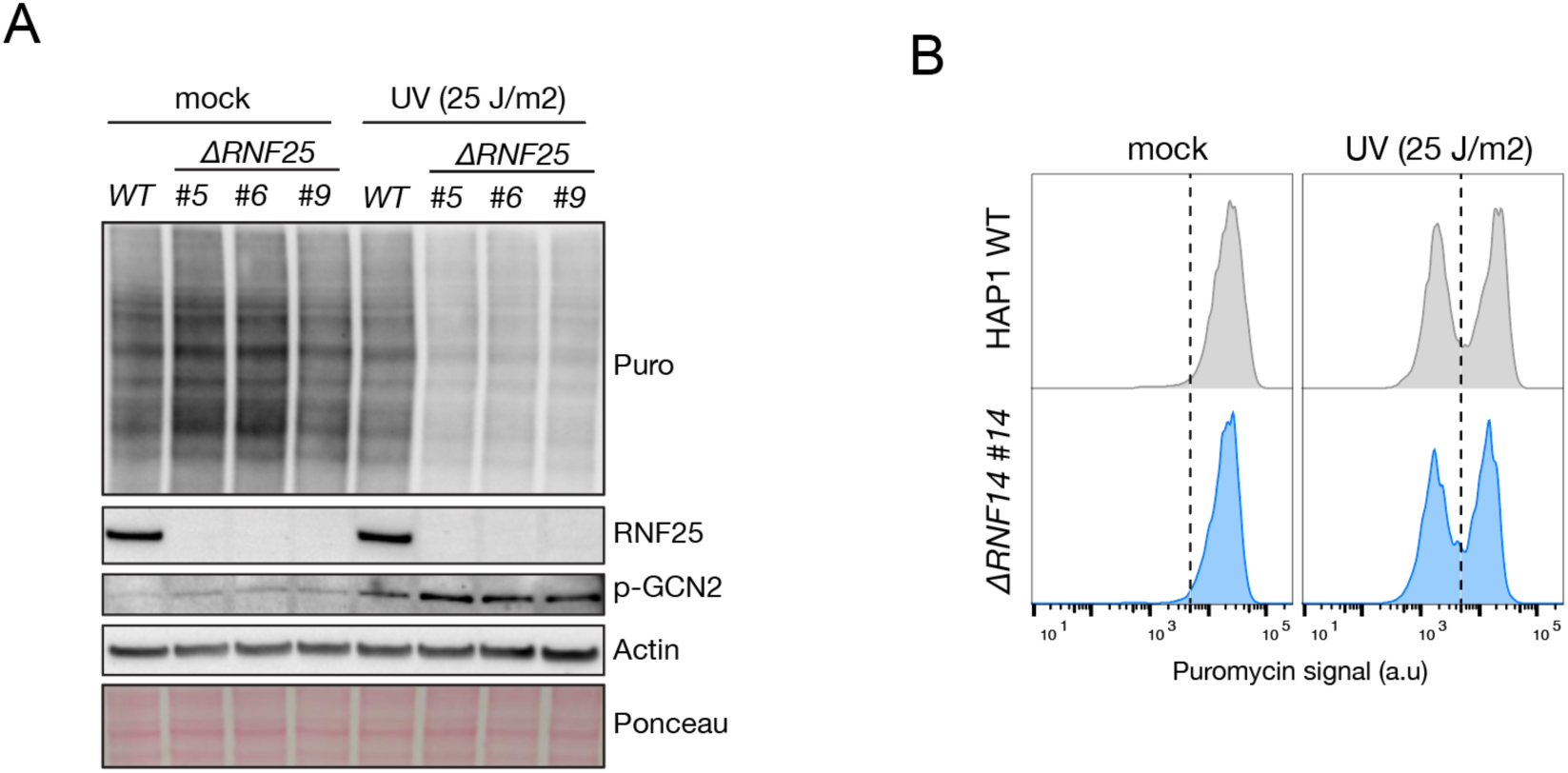
RNF25 promotes translation in response to UV radiation independently of RNF14. (**A**) Parental HAP1 and three ΔRNF25 clones were mock or UV-treated (25J/m^2^, 2hr recovery). Cells were then labeled with puromycin and subjected to immunoblot analysis using the indicated antibodies. (**B**) Flow cytometry analysis of puromycin incorporation after mock or UV treatment (25J/m^2^, 2 hr recovery) in parental HAP1 and ΔRNF14 cells.

**Fig. S3.**
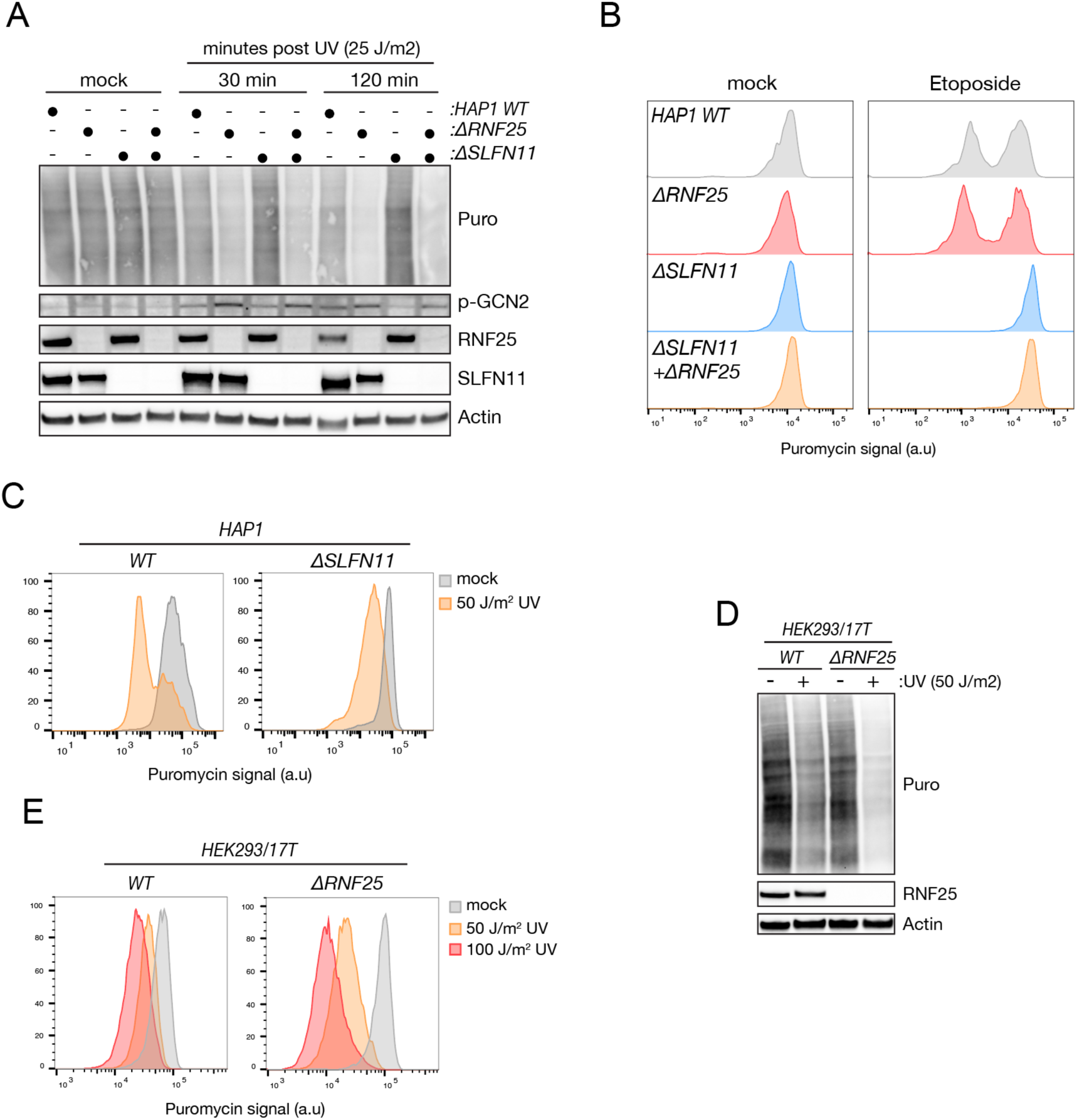
RNF25 is required for protein synthesis in response to RNA damage but not DNA damage. (**A**) Parental HAP1, ΔRNF25, ΔSLFN11 and ΔSLFN11ΔRNF25 cells were mock or UV-treated (25 J/m^2^). 30- and 120-mins following UV treatment, cells were labeled with puromycin and subjected to immunoblot analysis using the indicated antibodies. (**B**) As in (A) but cells were treated with Etoposide (5 μM, 3 hrs) and puromycin incorporation was analyzed by flow cytometry. (**C**) Flow cytometry puromycin incorporation assay in parental HAP1 and ΔSLFN11 cells treated with mock or high dose UV (50 J/m^2^, 2 hr recovery). (**D**) Parental HEK293/17T cells and ΔRNF25 cells were mock or UV-treated (50 J/m^2^, 2hr recovery). Cells were then labeled with puromycin and subjected to immunoblot analysis using the indicated antibodies. (**E**) As in (D) but cells were treated with 50J/m^2^ and 100J/m^2^ UV and puromycin incorporation analyzed by flow cytometry.

**Fig. S4.**
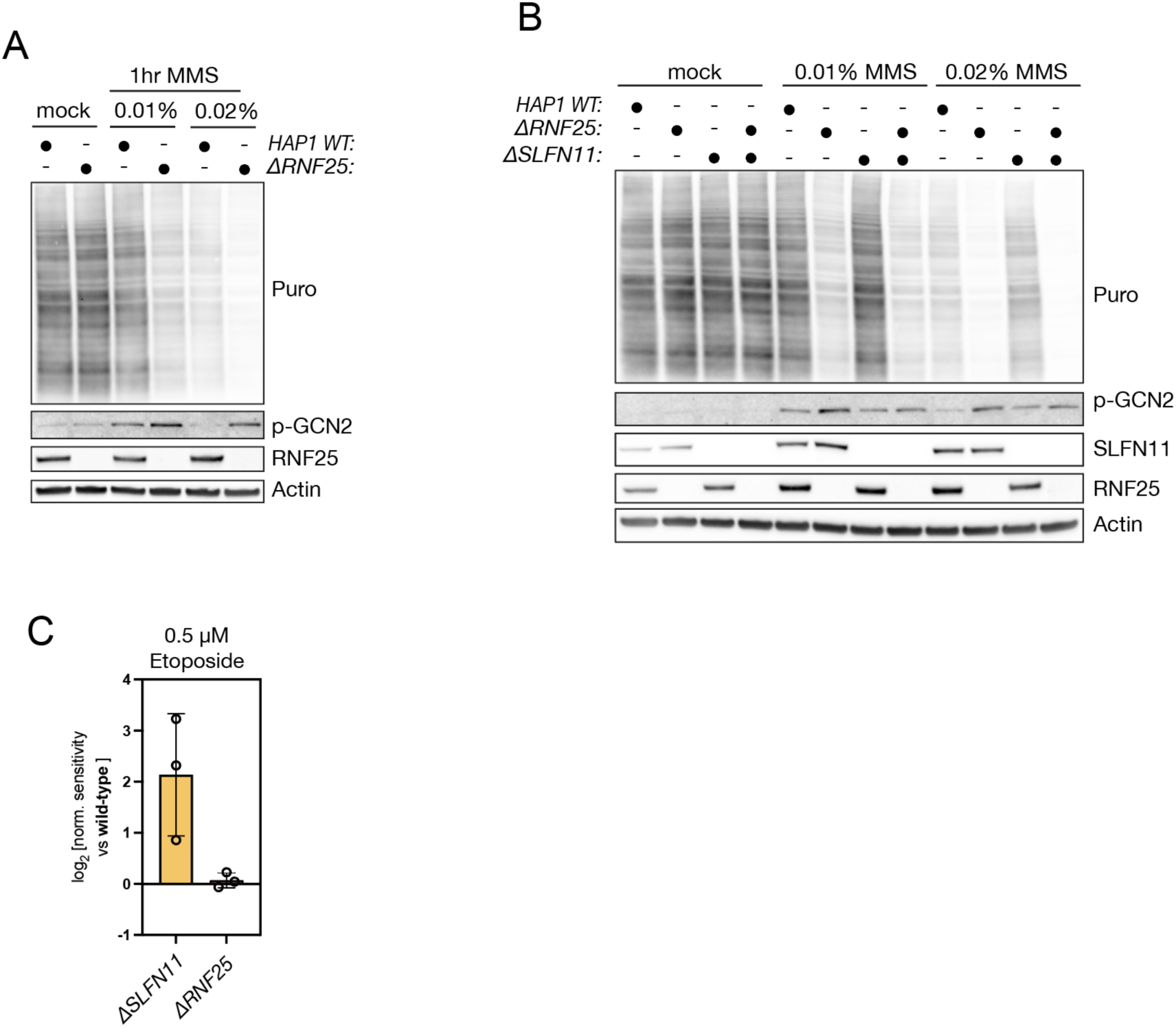
RNF25 supports protein synthesis and cell proliferation in response to RNA damage and not DNA damage. (**A**) Parental HAP1 and ΔRNF25 cells were treated with indicates concentrations of MMS for 1h. Cells were then labeled with puromycin and subjected to immunoblot analysis using the indicated antibodies. (**B**) As in (A) but ΔSLFN11 and ΔSLFN11ΔRNF25 cells were included. (**C**) Growth competition assay comparing proliferation of parental HAP1 cells to ΔSLFN11 and ΔRNF25 cells, respectively, in mock and etoposide-treated conditions (0.5 μM) for three days. Data represented as mean±SD of the normalized drug sensitivity ratio (log2). n = 3 independent biological replicates.

**Fig. S5.**
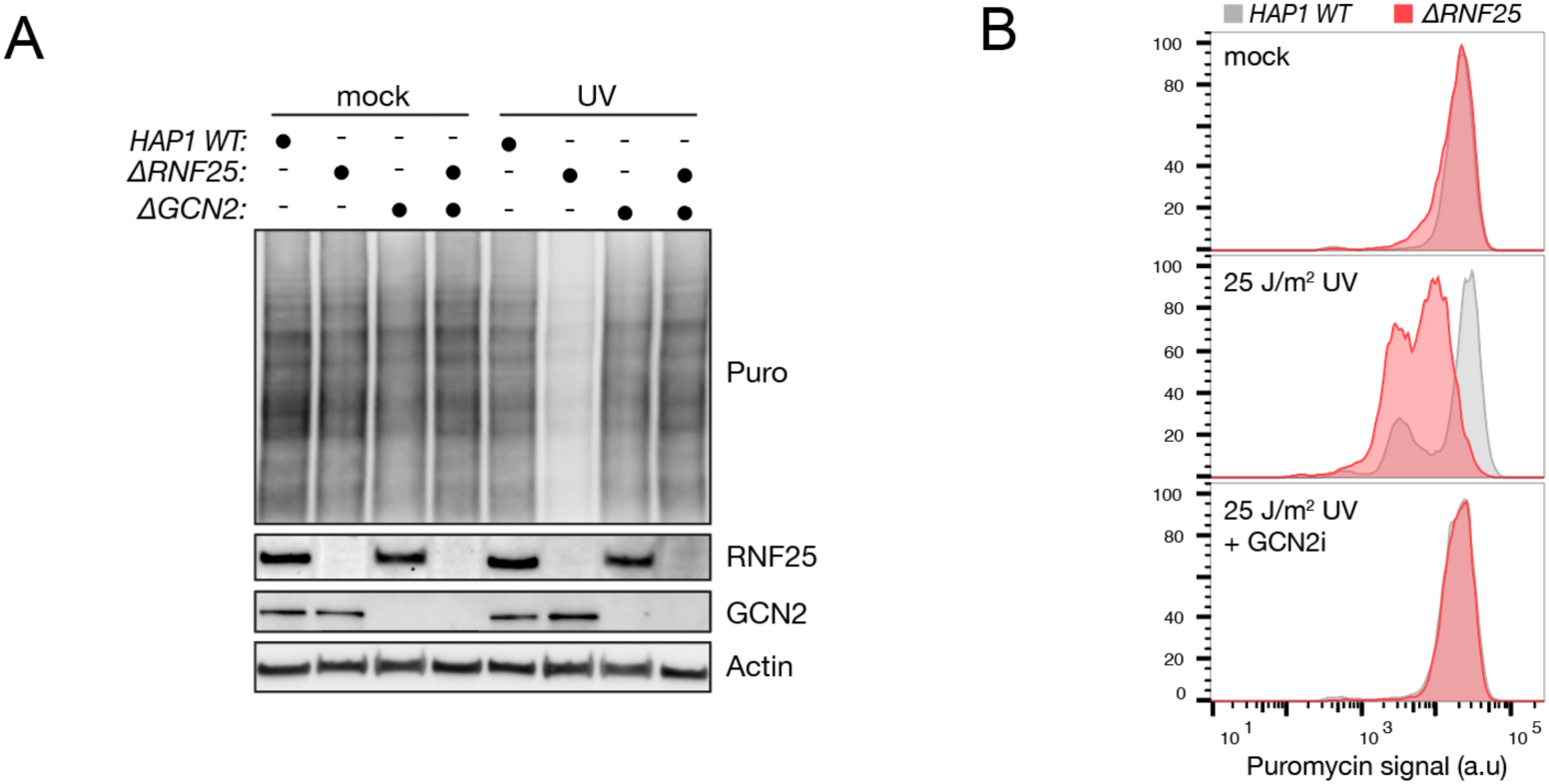
GCN2 loss or inhibition abrogates RNF25-dependent translation shutdown. (**A**) Parental HAP1, ΔRNF25, ΔGCN2 and ΔGCN2ΔRNF25 cells were mock or UV-treated (25J/m^2^, 2h recovery). Cells were then labeled with puromycin and subjected to immunoblot analysis using the indicated antibodies. (**B**) Flow cytometry analysis of puromycin incorporation after mock or UV treatment (25J/m^2^, 2 hr recovery) in parental HAP1 and ΔRNF25 cells in the presence or absence GCN2 inhibitor (GCN2i, A-92, 1 μM).

**Fig. S6.**
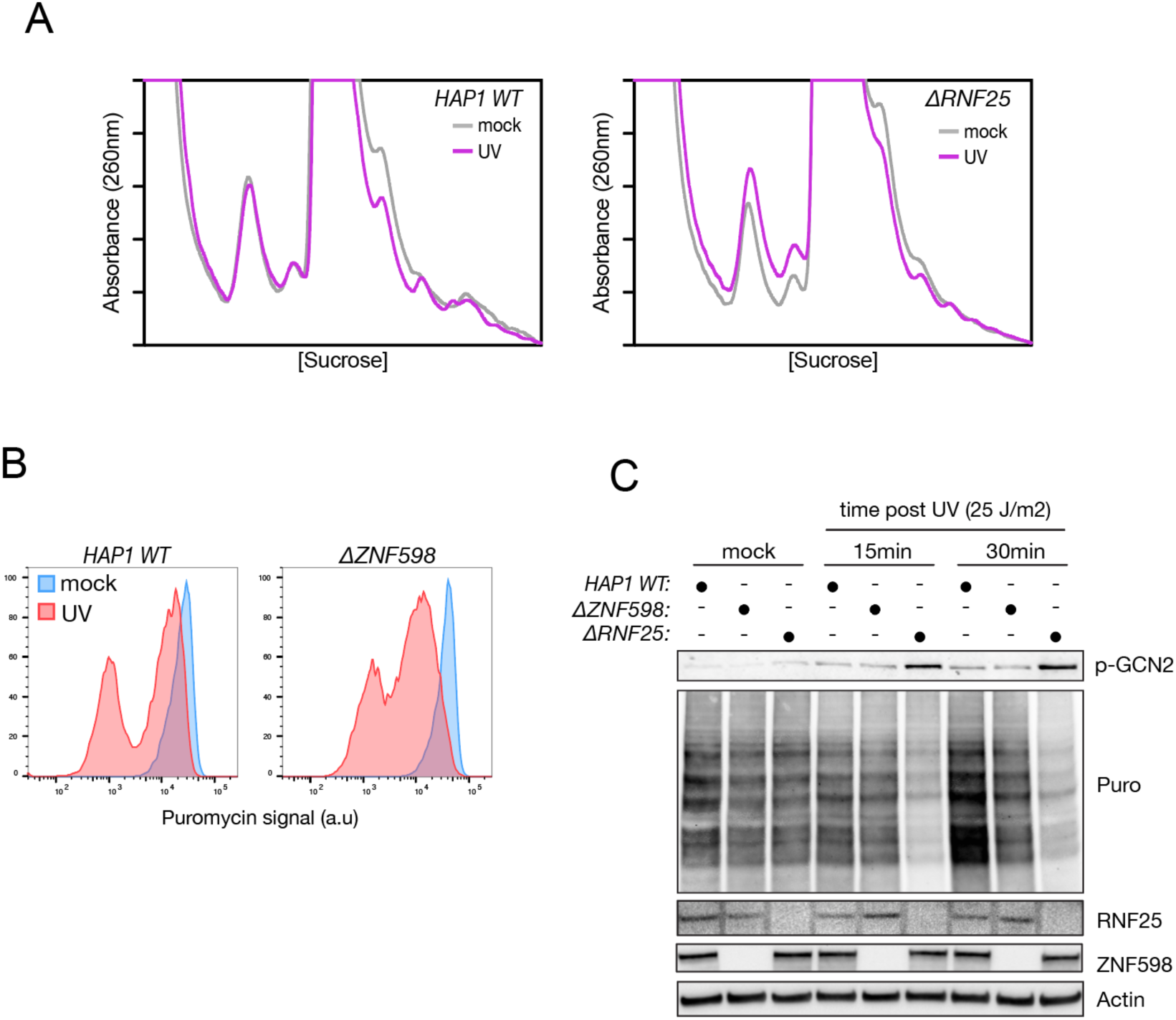
Aberrant accumulation of ribosome collisions are not underlying RNF25- dependent phenotypes in the UV response. (**A**) Disome profiles (MNase resistant) of parental HAP1 and ΔRNF25 cells in mock or UV-treated conditions (50J/m^2^, 30 min recovery). Cell lysates were digested with micrococcal nuclease (MNase) before passing through a sucrose gradient and continuously measuring absorbance at 260 nm. (**B**) Flow cytometry analysis of puromycin incorporation after mock or UV treatment (25J/m^2^, 2 hr recovery) in parental HAP1 and ZNF598- knockout (ΔZNF598) cells. (**C**) As in (B) but including ΔRNF25 cells and cells were subjected to immunoblot analysis to monitor bulk puromycin incorporation at indicated timepoints following UV treatment.

**Fig. S7.**
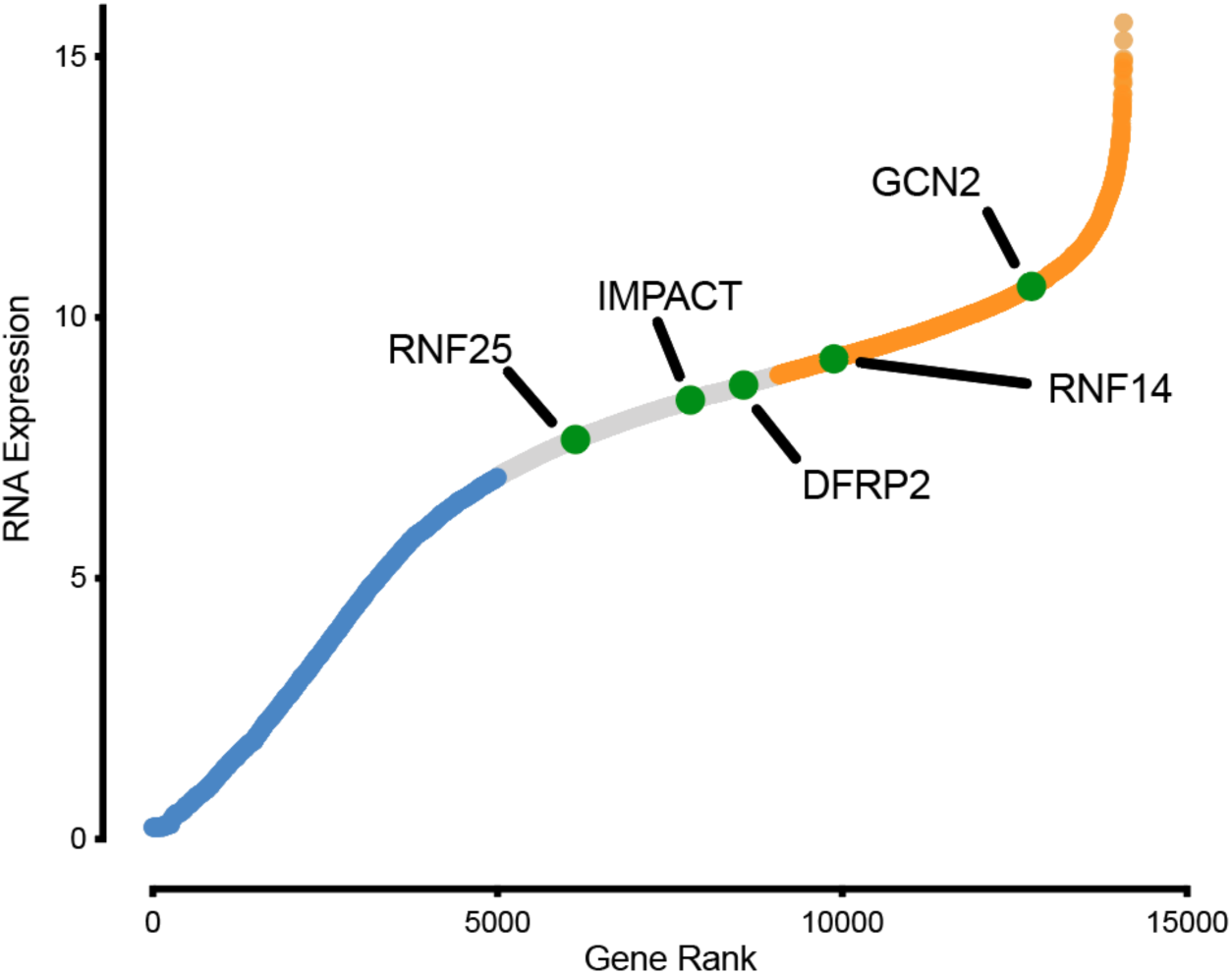
RNF25 is lower expressed in HAP1 cells compared to other reported RWD- containing proteins. (**A**) RNAseq data from parental HAP1 wild-type cells. Genes are ranked according to their transcript RNA expression, and the 5000 lowest and highest ranked genes are colored blue and orange, respectively, for visual purposes. Highlighted genes with validated RWD- domains are marked in green. Data acquired from Brockmann et al. 201C.^55^

## Methods

### Plasmids and cloning

Single guide RNAs (sgRNAs) were purchased as ssDNA oligos by IDT, annealed and then cloned into pX330 (Addgene #42230)) or pLentiCRISPRv2-Blast (Addgene #83480) using restriction enzymes BbsI or BsMBI, respectively. The following sgRNAs were used in this study: RNF25 (5’-GCGGGCCGGTGAAGATATGG-3’), RNF14 (5’- GCCCTGGCAAGTATTTACGA-3’), ZNF598 (5’- TAGAGCAGCGGTAGCACACC -3’), GCN2 (5’-TATATGTAAAAGTGGATTTG-3’). Expression vectors for RNF25-V5 WT and RNF25-V5 ΔRING (Δ134-165), as well as plasmids for fluorescent labeling with mEmerald or mCherry were acquired from addgene (pHR V5-RNF25_IRES-mCherry, Plasmid #198386; pHR V5-RNF25 dRING_IRES-mCherry, Plasmid #198387; pFUW mEmerald-P2A-PuroR, Plasmid #239566; pFUW mCherry-P2A-PuroR, Plasmid #239567). RNF25-V5 ΔRWD (Δ16- 125) was made from the pHR V5-RNF25_IRES-mCherry construct. In short, reverse PCR was done using primers designed to exclude the deleted regions, and with overhangs that would provide 20-40 base pair overlap 0-80 bp overlap between the ends. Gibson assembly was done using Gibson Assembly® Master Mix (NEB, #E2611), following their guidelines. The Gibson assembly product was subsequently transformed into One Shot™ Stbl3™ Chemically Competent E. coli (Thermofisher, C737303) and grown at 30 °C, followed by plasmid extraction.

### Cell culture and treatments

HAP1 cells were cultured in Iscove’s Modified Dulbecco’s Medium (IMDM) (#31980030, Thermo Fisher Scientific) supplemented with 10% heat inactivated fetal calf serum (FCS, Capricorn Scientific), GlutaMAX (Gibco), and penicillin streptomycin (P/S, Gibco) at 37°C in 5% CO2. HEK293T cells were cultured in Dulbecco’s modified Eagle’s medium (DMEM) (Thermo Fisher Scientific) supplemented with 10% FBS, GlutaMAX, and P/S at 37°C in 5% CO2.

To generate clonal knockout cell lines, parental cells were (co-)transfected with CRISPR/Cas9 vectors pX330 or pLentiCRISPRv2-Blast, containing the required sgRNA sequence, and a resistance cassette if needed (blasticidin), using Xfect (Takara Bio) according to the manufacturer’s instructions. Cells were then briefly selected with 40μg/ml blasticidin (ThermoFisher) for 48 hrs and allowed to recover for 24 hrs prior to seeding for clonal outgrowth. Gene status was monitored by sanger sequencing of genomic DNA and/or immunoblot analysis. All cell lines were further screened for ploidy status and only haploid clones were selected. Double knockout clones were generated similarly from a single knockout starting point.

Generation of stably expressing HAP1 cell lines was done polyclonally using lentiviral transduction. Briefly, lentivirus was produced in HEK293T cells transfected with pVSVG, pΔVPR, pAdvantage and a lentiviral expression plasmid using Lipofectamine 2000 according to the manufacturer’s protocol. The supernatant containing virus was collected two days after transfection and passed through a 40μm filter. The filtered viral supernatant was diluted in IMDM and applied to HAP1 cells supplemented with 8μg/ml protamine sulfate for 24 hours and recovered for 24 hours in fresh medium. Cells stably expressing fluorescent markers (pFUW mEmerald-P2A-PuroR, Plasmid #239566; pFUW mCherry-P2A-PuroR, Plasmid #239567) for the competition co-culture assay were selected using puromycin (10ug/ml) for 48h and assessed for fluorescence using live-cell flow cytometry. HAP1 cells transduced with pHR V5-RNF25_IRES-mCherry (#198386 or pHR V5-RNF25 dRING_IRES-mCherry, (#198386) were sorted in bulk based on mCherry expression and validated for RNF25-V5 expression using immunoblot analysis.

Reagents used in this study were DMSO (Sigma), etoposide (Sigma, E1383), puromycin (Thermo Fisher, 2 or 10 μg/ml), Methyl Methanesulfonate (MMS, 1-10 μg/ml, #129925- 25G, Sigma), 5-azacytidine (Sigma, 0.5 μg/ml), blasticidin (Thermo Fisher, 10 μg/ml), ISRIB (Sigma, 0.2 μM).

### Haploid genetic screens

Haploid genetic screens were performed as previously described with only small variations as briefly described below.^56^ A GFP-containing variant of gene-trap retrovirus was produced by transfecting HEK293T cells with pGT-GFP, pVSV-G, pΔVPR, and pAdvantage (Promega). Forty-eight hours post-transfection, viral supernatant was collected, passed through a 40 µm filter, and concentrated using centrifugal Amicon 100 kDa cut-off filters (#UFC9100, Merck), according to the manufacturer’s instructions.

Concentrated virus was kept at 4°C overnight. Fresh medium was then reapplied to the 293T cells, and virus collection and concentration were repeated after an additional 24 hours. The pooled concentrated virus was then immediately applied to HAP1 cells in the presence of 8 µg/ml protamine sulfate. After 24 hours, cells were recovered in fresh medium and cultured as described above for 8–10 days prior to use in genetic screens. For a genetic screen, cells were expanded to a total of 3-4 × 10⁹ cells and treated with UV irradiation (25 J/m^2^, 2 hr recovery), thapsigargin (4 μM, 1.5 hr), or sodium Arsenite (100 μM, 1 hr). Following treatment, cells were incubated with puromycin (2 µg/ml) for 15 minutes, after which the medium was removed, and cells were washed twice with PBS. Cells were then harvested by trypsinization, washed once with PBS, and subsequently fixed, permeabilized, and blocked as described in the *Flow Cytometry* methods section. Cells were stained in 30 mL FACS buffer with a primary antibody against puromycin coupled to AlexaFluor-647 and DAPI (1 ug/ml, #D9542, Sigma-Aldrich), passed through a 40 µm filter, and sorted on a FACSAria Fusion cell sorter (BD Biosciences). Haploid G1 cells were gated based on DAPI staining and >1 × 10⁷ cells were collected for both the high and low populations, representing the 5% of cells with the highest and lowest puromycin fluorescence signal, respectively.

Genomic DNA isolation from sorted cells was done using the QIAamp DNA Mini Kit (Qiagen, 51306). Sequencing library preparation, and sequencing analysis were performed as previously described.^57^ Briefly, Genomic DNA was subjected to linear amplification–mediated PCR (LAM-PCR) using dual-biotinylated primers and AccuPrime Taq polymerase. Reactions were amplified for 120 linear cycles (94°C, 58°C, 68°C). Biotinylated LAM products were captured using streptavidin-coated magnetic beads (brand) and washed extensively. A DNA linker was ligated to bead-bound LAM PCR products using CircLigase II ssDNA ligase (Biosearch Technologies). Following ligation, samples were incubated at at 80°C for 10 min to heat inactivate the ligase and washed extensively. Libraries were then amplified by adapter PCR using multiplex primers. Libraries were assessed by agarose gel electrophoresis, purified using PCR purification kit (QIAgen), quantified by Qubit, and analyzed by TapeStation using a D5000 prior to sample pooling and sequencing. Amplified libraries were sequenced on a NextSeq2000 using a P2-100 sequencing kit with a read length of 75 bp (single end read). Sequencing reads were aligned to the GRCh38 reference genome allowing one mismatch and assigned to protein-coding regions. A mutational index (MI) was calculated by comparing the number of insertions in the sense direction in the high population to that of the low populations using a two-sided Fisher’s exact test to identify significant differences (FDR- corrected *p* < 0.05). Fishtail scatterplots were generated using GraphPad Prism by plotting the MI of each gene (log2, y-axis) to the total number of unique insertions of the gene identified in the two populations (log10, x-axis).

### Immunoblotting analysis

Cells for whole cell extract immunoblot analysis were washed in PBS and lysed directly in 2x LDS sample buffer (Bolt^TM^, Invitrogen) supplemented with 100μM DTT, sonicated using a stick sonicator, and incubated for 10 minutes at 95 °C. Proteins were then resolved by SDS-PAGE, transferred onto nitrocellulose membranes (0.45 μm pore size) and blocked in 5% milk in TBST (TBS with 0.1% Tween) for at least 30 min. Membranes were then incubated with primary antibody (diluted in 5% milk TBST) overnight at 4 °C, washed in TBST and incubated with the corresponding secondary antibody coupled to HRP (Thermo Fisher Scientific) for 1 hour at RT. Following thorough washing in TBST, SuperSignal West Pico PLUS (Thermo Fisher Scientific) was used as HRP substrate according to manufacturer’s instructions, and membranes were imaged on a Universal Hood II Gel Doc system (Bio-Rad). All immunoblotting experiments were performed at least twice, and representative data are shown.

To assess puromycin incorporation by immunoblot, cells were incubated with puromycin (2μg/ml) for 10 minutes before processing as described above.

Antibodies used for immunoblot in this study were anti-puromycin (Millipore, MABE343,1:5000), anti-RNF25 (abcam, ab140514,1:1000), anti-pGCN2 Thr899 (Cell Signaling, 94668S, 1:1000), anti-RNF14/Ara54 (santa cruz, sc-376701, 1:1000), anti- SLFN11 (Santa Cruz, 515071, 1:1000), anti-GCN2 (Cell Signaling Technology, 3392, 1:1000), anti-p-JNK (Cell Signaling Technology,4668, 1:1000), anti-actin(#ab123034, clone 2A3, abcam). Anti-V5 (abcam, ab 27671, 1:1000).

### Flow cytometry

Flow cytometry analysis to assess puromycin incorporation was performed as follows. Cells were seeded 24 h prior to treatment and exposed to the indicated conditions for the specified durations. To label nascent protein synthesis, puromycin (2 µg/ml) was added directly to the culture medium 10 min prior to cell harvesting. Cells were subsequently washed twice with PBS, trypsinized, and collected directly into FACS buffer (PBS supplemented with 5% FBS). Cell suspensions were washed once with FACS buffer and fixed using Fix Buffer I (*BD Biosciences*) for 10 min at 37 °C. Following fixation, cells were washed twice with FACS buffer and stored at 4 °C until further processing.

Cells were permeabilized using Perm Buffer III (*BD Biosciences*) for 30 min on ice and washed in FACS buffer. Samples were then barcoded to enable pooling of multiple conditions using fluorescent NHS Ester barcoding dyes (Alexa Fluor 488, Alexa Fluor 800, *ThermoFisher*) at concentrations of 0.4, 2, and 5 µg ml⁻¹ diluted in 70% ethanol.^58^ Barcoding was performed for 15 min at room temperature, followed by two washes with FACS buffer. Barcoded samples were subsequently pooled appropriately and incubated with DAPI (1 ug/ml, #D9542, Sigma-Aldrich) and an Alexa Fluor 647– conjugated anti-puromycin antibody (Sigma) for 90 min at room temperature. Cells were washed twice with FACS buffer and analyzed on a FACSAria Fusion (BD Biosciences). Antibodies used for flow cytometry in this study were: Alexa Fluor® 647 Anti-Puromycin (Sigma, MABE343-AF647).

For flow cytometry analysis of fluorescent markers in live-cells, cells were trypsinized and directly resuspended in FACS buffer (PBS with 5% FBS) and kept on ice until analyzed on a FACSAria Fusion (BD Biosciences). Data was exported and processed using FlowJo software.

### Growth Competition Assay

Growth competition assays were performed using co-cultures of fluorescently labeled cell lines. Parental HAP1 and derived cell lines stably expressing mEmerald- or mCherry- were mixed in the required comparisons at a 1:1 ratio and seeded into 24-well plates, with each condition plated in technical triplicates. Cells were allowed to adhere for 24 h and left untreated (normalization control) or exposed to the indicated drugs: methyl methanesulfonate (MMS; 1.5 or 2 µM), etoposide (0.5 µM), ISRIB (0.2 µM), or 5-azacytidine (5-aza; 2 µM). Treatments were maintained for 3 or 5 days as indicated. At the end of the treatment period, cells were trypsinized and collected in FACS Buffer (PBS with 5% FBS) and the relative proportions of mEmerald- and mCherry-positive cells were quantified by flow cytometry. All experiments were performed in technical triplicates and repeated in at least three independent biological replicates.

The normalized sensitivity ratio score was calculated as the ratio of the two fluorescent populations (A:B, e.g. wild-type vs knock-out) in treated samples, normalized to the corresponding ratio in the untreated control. Data are presented as log₂ fold change of the treated-to-untreated ratio, such that 0 indicates no differential growth effect between genotypes wheras positive and negative values indicate a relative growth advantage or disadvantage, respectively, of the compared genotype.

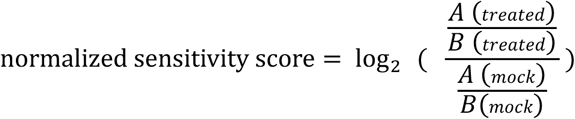

### Disome analysis by polysome profiling

Cells were plated the day before and treated with indicated UV irradiation dose or a mock treatment. 30 mins following UV treatment, cells were scraped in ice cold PBS containing 100 μg/ml cycloheximide and lysates were prepared using 20 mM HEPES pH 7.5, 100 mM NaCl, 5 mM MgCl2, 100 μg/ml digitonin, 100 μg/ml cycloheximide, 1X protease inhibitor cocktail (Sigma, #P2714) and 1μM dithiothreitol (DTT). Extracts were pushed 4 times through a 27G needle and incubated on ice for 5 minutes prior to centrifugation at 17,000 g for 5 minutes at 4 °C. The amount of RNA in each lysate was quantified with Nanodrop and equivalent amounts were used for MNAse digestion (disome analysis) and loading on gradients. 1mM CaCl2 was added to the lysates together with 500 U micrococcal nuclease (MNase) (New England Biolabs, #M0247) and samples were incubated for 30 minutes at 22 °C ^32^ . Digestion was stopped with 2mM EGTA and samples were centrifuged at 17,000 g for 5 minutes at 4 °C. Lysates were resolved on a 15-50% sucrose gradient by centrifugation at 38,000 rpm for 2 hours at 4 °C in a Sorvall TH64.1 rotor. Gradients were analyzed using a Biocomp density gradient fractionation system with continuous monitoring of the absorbance at 260 nm.

